# Swapping Birth and Death: Symmetries and Transformations in Phylodynamic Models

**DOI:** 10.1101/494583

**Authors:** Tanja Stadler, Mike Steel

## Abstract

Stochastic birth–death models provide the foundation for studying and simulating evolutionary trees in phylodynamics. A curious feature of such models is that they exhibit fundamental symmetries when the birth and death rates are interchanged. In this paper, we explain and formally derive these transformational symmetries. We also show that these transformational symmetries (encoded in algebraic identities) are preserved even when taxa at the present are sampled with some probability. However, these extended symmetries require the death rate parameter to sometimes take a negative value. In the last part of this paper, we describe the relevance of these transformations and their application to computational phylodynamics, particularly to maximum likelihood and Bayesian inference methods, as well as to model selection. Phylodynamics, phylogenetics, speciation-extinction models, birth-death models, algebraic symmetries, maximum likelihood, Bayesian inference

## 1 Introduction

Linear birth–death models play a pivotal role in phylodynamics. These stochastic models provide a prior distribution on evolutionary trees (both the shape and edge length distribution) for Bayesian inference methods [21, 18]. Moreover, these models allow biologists to estimate key parameters of macroevolution (such as speciation rates corresponding to birth rates, and extinction rates corresponding to death rates) from reconstructed phylogenetic trees which were dated by fossil (or other time-sampled) evidence [10].

The study of such models dates back to some classical papers from the early to mid-20th century [22, 5, 6], and the application of these models to phylogenetics and phylodynamics flourished from the 1990s onwards [10, 11]. Further in-depth mathematical analysis [1, 8, 2, 9, 7] has extended our understanding of the properties of these models and extensions that allow more complex processes of birth and death.

In this paper, we identify and explore curious symmetries in fundamental birth–death model probability distributions when the birth and death rates (*λ* and *μ*) are swapped. We will start the paper by providing an intuitive account of this symmetry that seems at first a little surprising. We extend this to the more general setting where a third parameter is introduced — the sampling probability *ρ* of taxa sampled at the present — and show how analogous symmetries can be derived by a transformation that reduces these three parameters to just two (*λ*′, *μ*′). One can view these as ‘corrected’ birth and death rates, except for the caveat that this new death rate *μ*′ can now take negative values. A major advantage of working with the transformed pair of parameters (*λ*′, *μ*′) is that it captures the correct dimensionality of the process (namely 2), thereby avoiding the inherent redundancy present in the 3-dimensional parameterization that uses the triple (*λ, μ, ρ*). This viewpoint has implications for phylogenetic and phylodynamic inferences, both in the maximum likelihood and Bayesian settings, and we explore these implications in the latter part of our paper.

## 2 Birth–death symmetries

Consider a phylogenetic tree that evolves from a single ancestral taxon according to a birth–death process, with a constant birth rate *λ* ≥ 0 and a constant death rate *μ* ≥ 0. Suppose that at some time point in the tree, there are *n* taxa present. Let *P_n,m_*(*t* | *λ, μ*) be the probability that at time *t* later, there will be *m* taxa present. These transition probabilities are classical and provide a foundation for phylodynamic models. However, the starting point for this paper is the following curious symmetry (communicated to the first author by Joseph Felsenstein):

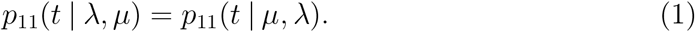

This equation states the surprising result that the probability of one individual having one surviving descendant after time *t* remains the same if we swap the birth rate (*λ*) and the death rate (*μ*). Thus a process with a birth rate of, say, 100 and a death rate of, say, 1 — a scenario with a very fast-growing population — has the same probability of having one surviving descendant as a process with a birth rate of 1 and a death rate of 100, a scenario where we know that the process eventually leads to extinction.

We will see that identities such as Eqn. (1) fall out from an algebraic analysis of birth–death models (provided later). Our aim in the meantime is to provide an intuitively transparent (but still rigorous) argument for Eqn. (1), as well as the following more general identity, namely that the probability of *n* individuals having *n* surviving descendants after time *t* is the same for birth rate *λ* and death rate *μ* or birth rate *μ* and death rate *λ*, for all *n* ≥ 1.

### Proposition 2.1.

*For any non-negative value of λ, μ and any value of n* ≥ 1:

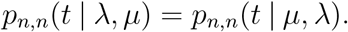

We now provide a direct and intuitively-transparent proof of Proposition 2.1. We deal in detail with the case *n* =1 (i.e. Eqn. (1)); however, the result for *n* ≥ 1 follows by essentially applying the same idea. We start a birth–death process with one individual. The waiting time between ‘events’ (a birth event or death event) is exp(*n*(*λ* + *μ*)), where *n* is the number of individuals at the considered time point. Let 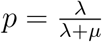, and consider two different scenarios (one proceeds forward in time, the other backward):

- Scenario 1: The process starts at time 0 and is stopped at time *t* > 0. At an event, with probability *p*, we add an individual and, with probability 1 − *p*. we remove an individual. Scenario 1 is a classic forward-in-time birth–death process.
- Scenario 2: The process starts at time *t* > 0 and is stopped at time 0. At an event, with probability 1 − *p* we add an individual and, with probability *p*, we remove an individual. Scenario 2 is a birth–death process in reversed time with the birth and death rates being interchanged compared with Scenario 1.

Intuitively, the result of the time-reversed process with birth and death being interchanged is analogous to the forward-in-time process. However, we justify this intuition by a formal argument showing that the probability of observing one individual after time *t* is the same under Scenario 1 and Scenario 2.

Consider some population size trajectory *X* that starts at time 0 with one individual and ends with one individual after time *t*. At each event, *X* can grow or decrease by one. Let the number of growth events be *k*, which therefore also equals the number of death events. Denote the time of these 2*k* events by *t*_1_, *t*_2_,… *t*_2*k*_, and define *t*_0_ = 0 and *t*_2*k*+1_ = *t*. See Figure 2 for an example with *k* = 2.

**Figure 1:**
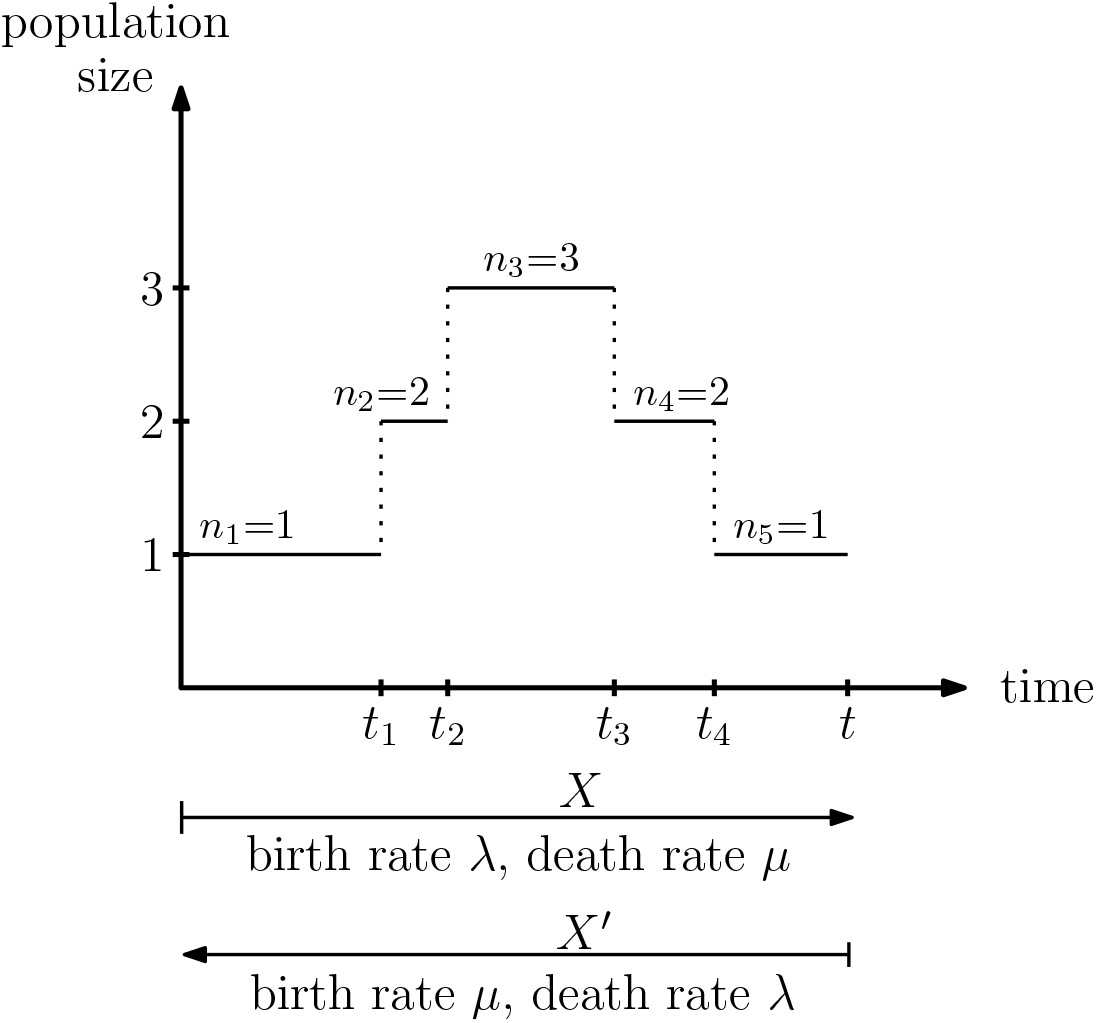
The forward-in-time birth–death process with realization *X* and the equivalent time-reversed process with interchanged rates and realization *X*′.

**Figure 2:**
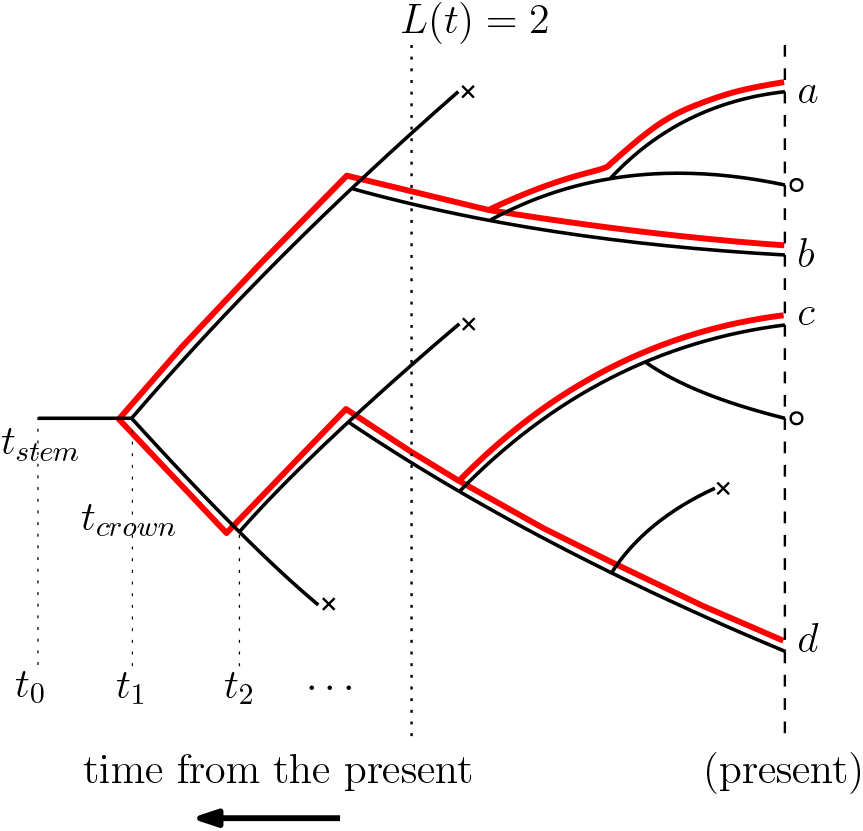
A phylogenetic tree 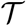 that evolves under under a birth–death process with rates *λ, μ* and with sampling at the present with probability *ρ*. Lineages ending in a death (extinction) are marked by × whereas lineages at the present that are not sampled are marked by o. The reconstructed tree on the sampled extant taxa is shown in red.

The probability density of *X* under Scenario 1, *L*_1_(*X*), is a product of the probability for the birth events, *p^k^*, for the death events (1 − *p*)^*k*^, and the waiting times between events, 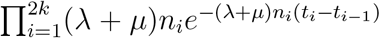, where *n_i_* is the number of individuals prior to the event at time *t_i_*. Finally, the term *e*^−(*λ*+*μ*)(*t*−*t*_2*k*_^) stipulates that no subsequent event happens after the event at time *t*_2*k*_. In summary, the probability density of *X* under Scenario 1 for *k* > 0 is:

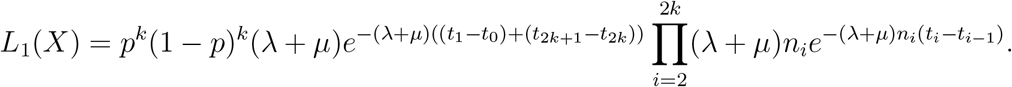

For *k* = 0, we have

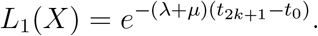

Now we reverse time in the realization *X* and call it *X*′. Thus, *X*′ starts where *X* ends, and *X*′ ends where *X* starts. The probability density of *X*′ under Scenario 2 is then *L*_2_(*X*′). We establish *L*_2_(*X*′) analogous to the procedure above, with the birth events in *X* being death events in *X*′ and vice versa. Thus, the same *p* and (1 − *p*) factors are multiplied when calculating the probability density of *X*′ under Scenario 2, compared to the probability density of *X* under Scenario 1. Furthermore, the waiting time contributions are the same for Scenario 1 and Scenario 2, and thus *L*_1_(*X*) = *L*_2_(*X*′).

Note that *p*_1_(*t* | *λ, μ*) is the integral over all realizations *X* under Scenario 1, 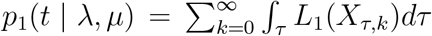, where *X_τ,k_* is a realization with *k* birth events according to an event time vector *τ* = (*t*_1_, *t*_2_,…, *t*_2*k*_).

Analogously, 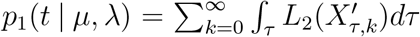. Since 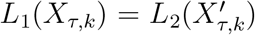, each component in this integration has the same probability density and thus we have *p*_1_(*t* | *λ, μ*) = *p*_1_(*t* | *μ,λ*).

One can directly extend this argument to establish Proposition 2.1 for any value of *n* ≥ 1 by considering the associated forward-in-time and backward-in-time processes. However, as we will derive this equation later from an algebraic identity, we do not describe this further here.

## 3 General symmetries under incomplete sampling

We continue to study a birth–death model with constant and non-negative birth and death rates *λ* and *μ*. However, we now allow each of the individuals present at time *t* to be sampled (independently) with probability *ρ* ∈ (0, 1].

Let us first suppose that we start with one individual at time *t*_0_, and let *p_i_*(*t* | *λ, μ, ρ*) be the probability that *i* sampled descendants are observed (i.e. extant and sampled) at time *t*_0_ + *t*. Exact expressions for *p_i_*(*t*) = *p_i_*(*t* | *λ, μ, ρ*) are provided by the following formulae (all proofs of theorems and corollaries are found in the Appendix). Let

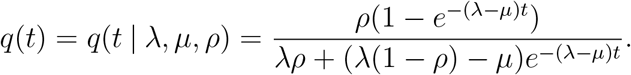

### Theorem 3.1.

*For λ* ≠ *μ, we have:*

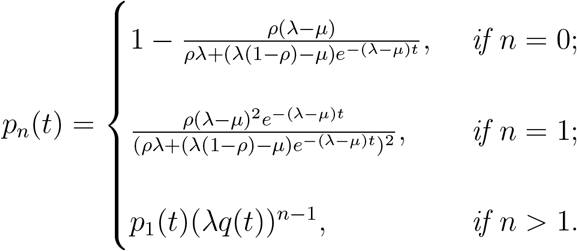

*Note that λq*(*t*) = *p*_2_(*t*)/*p*_1_(*t*), and for *ρ* = 1 and *μ* > 0, we have 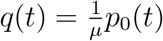.

### Corollary 3.2.

*For the critical case* 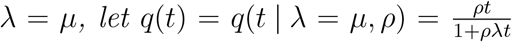. *We then have:*

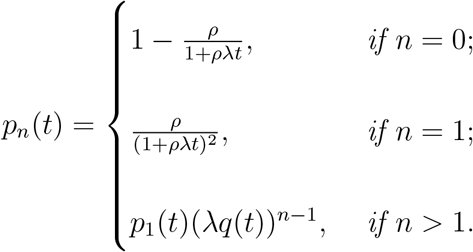

We investigate the expressions for *p_i_*(*t* | *λ, μ, ρ*) in detail, and identify symmetries with respect to *λ* and *μ*. We begin with the case where all extant taxa are sampled.

### Theorem 3.3.

*In the case of complete sampling (i.e. ρ* = 1*), we have:*

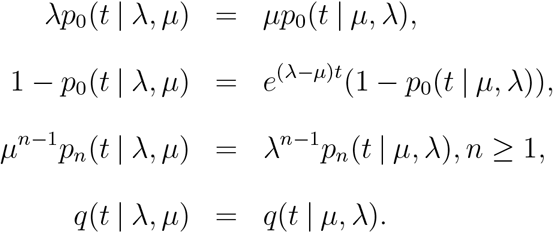

Next, consider the probability *p_n,m_*(*t* | *λ, μ*) of having *m* individuals at time *t*, given that *n* are present at time 0 (as described in the introduction).

### Corollary 3.4.

*For m* ≥ *n* ≥ 1, *we have: μ*^*m*−*n*^*p_n,m_*(*t* | *λ, μ*) = *λ*^*m*−*n*^*p_n,m_*(*t* | *μ, λ*). *In particular, p_n,n_*(*t* | *λ, μ*) = *p_n,n_*(*t* | *μ, λ*), *for all n* ≥ 1 *(as in Proposition 2.1). For* 0 ≤ *m* < *n, we have: λ*^*n*−*m*^*p_n,m_*(*t* | *λ, μ*) = *μ^n-m^p_n,m_*(*t* | *μ, λ*).

### 3.1 Negative ‘death rates’ in the case of incomplete sampling

We now investigate the case *ρ* ≤ 1. We introduce two new variables *λ*′ and *μ*′, which will play a key role in the remainder of the paper. They are defined by *λ, μ* and *ρ* according to the following transformation:

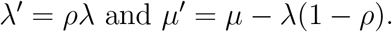

Note that when *ρ* = 1, we have *λ*′ = *λ*. Further, for all vales of *ρ* we have *λ*′ – *μ*′ = *λ* – *μ* (thus *λ*′ ≠ *μ*′ if and only if *λ* ≠ *μ*). Note also that *μ*′ < 0 is entirely possible (for example, when *λ* = 4*μ* and *ρ* = 0.5, we obtain *μ*′ = –*μ*). In this case, *μ*′ can not easily be viewed as a death rate (nor as a birth rate); however, allowing *μ*′ to take any real value (positive or negative) means that all parameter triplets (*λ, μ, ρ*) have a transformation to (*λ′, μ′*).

The following lemma is straightforward to verify using simple algebra.

#### Lemma 3.5.

*For all λ, μ* ≥ 0 *and* 0 < *ρ* < 1, *the four functions*

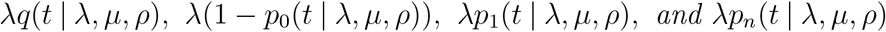

*can be written as functions of only two parameters* (*λ*′ *and μ*’) *when λ* ≠ *μ (rather than the three parameters λ, μ, ρ). When λ* = *μ, these four functions can be written as functions of the single parameter λ*′.

In order to investigate symmetries, we define the following functions, which only depend on *λ*′, *μ*′, and *t* (rather than the four parameters *λ, μ, ρ* and *t*). Let:

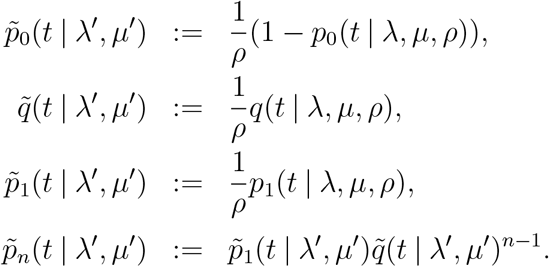

For *λ* ≠ *μ*, these equations are,

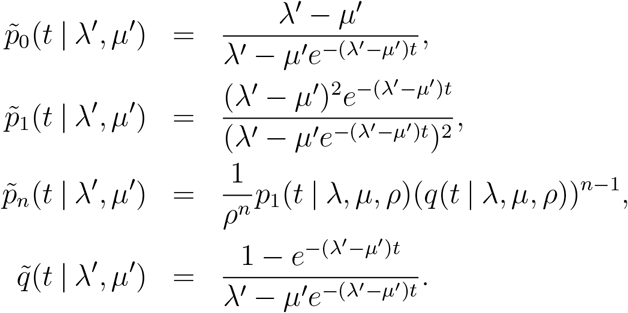

In particular, we have: 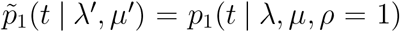. This leads to the following symmetries with respect to *λ*′ and *μ*′.

#### Theorem 3.6.

*For μ*′ ≥ 0, *the following symmetries hold:*

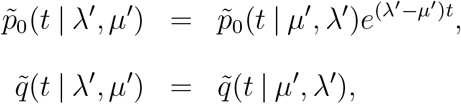

*and for all n* ≥ 1:

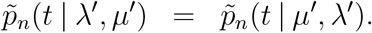

## 4 Tree probability densities

Let 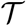 be a phylogenetic tree generated by a birth–death process starting with one taxon and being stopped after time *t*_0_. Each individual alive after time *t*_0_ is sampled with probability *ρ*. In this tree, all extinct lineages are pruned, and only the lineages leading to the sampled tips are kept. Such a tree is also called the *reconstructed tree* [10], as indicated by the red lines in Fig. 2. Let this tree have *n* sampled tips and the branching times *t*_1_ > *t*_2_,… > *t*_*n*−1_, where time is measured from the present time 0. Let *L*(*t*) be the number of co-existing lineages of tree 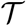 at time *t* (see Fig. 2).

Let 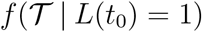 be the probability density of the tree 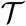, and let 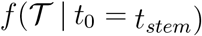 be the probability density of the tree 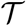, given that at least one individual is sampled at present. Thus *t*_0_ is the stem age (*t_stem_*) of the process. For *ρ* =1, this corresponds to conditioning on non-extinction of the process. Let 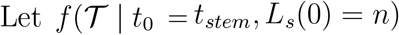 denote the probability density of the tree 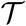, given that we sample exactly *n* tips at present (denoted by *L_s_*(0) = *n*).

The tree 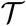 in these formulations was a tree starting with one individual, leading to two lineages at time *t*_1_ in the past. Alternatively, a tree 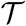 may start with two lineages at time *t*_1_ ago; the probability of such a tree is 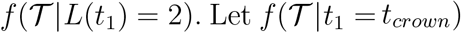 be the probability density of the tree 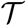 conditioning on sampling at least one descendant individual from both initial lineages. Note that when conditioning on sampling, the time *t*_1_ is the crown age of the clade (*t_crown_*). Furthermore, let 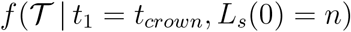 be the probability density of the tree 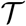 conditioned on sampling exactly *n* tips at present. Finally, in the setting where *t*_0_ is chosen uniformly at random from (0, ∞), then a tree 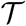 conditioned on *n* tips and integrated over all possible *t*_0_ has probability density 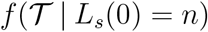.

In what follows, we assume *λ* > 0 and thus *λ*′ > 0; otherwise, we cannot obtain a tree with *n* > 1.

### Theorem 4.1.

*The tree probability densities can be expressed as functions of p*_0_(*t* | *λ, μ, ρ*), *p*_1_ (*t* | *λ, μ, ρ*) *and q*(*t* | *λ, μ, ρ*), *or* 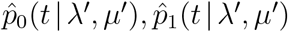 *and* 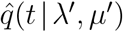. *Omitting the parameters λ, μ, ρ, λ′ and μ′ in these functions for easier reading, the expressions are given in the following table:*

| Tree probability densities | $(\lambda, \mu, \rho)$ – parameters | $(\lambda', \mu')$ – parameters |
| --- | --- | --- |
| Unconditioned |  |  |
| $f(\mathcal{T} \mid L(t_0) = 1)$ | $p_1(t_0) \prod_{i=1}^{n-1} \lambda p_1(t_i)$ | $\rho \tilde{p}_1(t_0) \prod_{i=1}^{n-1} \lambda' \tilde{p}_1(t_i)$ |
| $f(\mathcal{T} \mid L(t_1) = 2)$ | $p_1(t_1)^2 \prod_{i=2}^{n-1} \lambda p_1(t_i)$ | $(\rho \tilde{p}_1(t_0))^2 \prod_{i=2}^{n-1} \lambda' \tilde{p}_1(t_i)$ |
| Conditioned |  |  |
| $f(\mathcal{T} \mid t_0 = t_{stem})$ | $\frac{p_1(t_0)}{1-p_0(t_0)} \prod_{i=1}^{n-1} \lambda p_1(t_i)$ | $\frac{\tilde{p}_1(t_0)}{\tilde{p}_0(t_0)} \prod_{i=1}^{n-1} \lambda' \tilde{p}_1(t_i)$ |
| $f(\mathcal{T} \mid t_1 = t_{crown})$ | $\left( \frac{p_1(t_1)}{1-p_0(t_1)} \right)^2 \prod_{i=2}^{n-1} \lambda p_1(t_i)$ | $\left( \frac{\tilde{p}_1(t_0)}{\tilde{p}_0(t_0)} \right)^2 \prod_{i=2}^{n-1} \lambda' \tilde{p}_1(t_i)$ |
| $f(\mathcal{T} \mid L_s(0) = n)$ | $n \frac{p_1(t_1)}{1-p_0(t_1)} \prod_{i=1}^{n-1} \lambda p_1(t_i)$ | $n \frac{\tilde{p}_1(t_0)}{\tilde{p}_0(t_0)} \prod_{i=1}^{n-1} \lambda' \tilde{p}_1(t_i)$ |
| $f(\mathcal{T} \mid t_0 = t_{stem}, L_s(0) = n)$ | $\prod_{i=1}^{n-1} \frac{p_1(t_i)}{q(t_0)}$ | $\prod_{i=1}^{n-1} \frac{\tilde{p}_1(t_i)}{\tilde{q}(t_0)}$ |
| $f(\mathcal{T} \mid t_1 = t_{crown}, L_s(0) = n)$ | $\frac{1}{(n-1)} \prod_{i=2}^{n-1} \frac{p_1(t_i)}{q(t_0)}$ | $\frac{1}{(n-1)} \prod_{i=2}^{n-1} \frac{\tilde{p}_1(t_i)}{\tilde{q}(t_0)}$ |

*We note that the expressions in the middle column have been presented in [13] [Eq. 1-7], highlighting that* 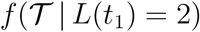 *goes back to [20] for ρ* = 1, 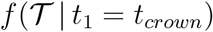 *to [10], and* 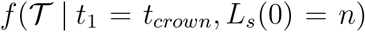 *to [21] (both for ρ* ∈ (0, 1]). *Furthermore*, the probability density 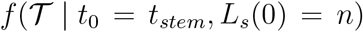 *for ρ* = 1 *is described in [4] and in earlier work by [12]. The idea of parameter transformation (right column) has been introduced for* 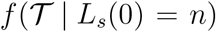 *in [14]*.

### Remark 4.2.

Only the expressions for the unconditioned tree probability densities (i.e. the equations not conditioning on observing at least one sample) depend on all three parameters *λ, μ* and *ρ*. The remaining five expressions (the conditioned tree probability densities) only depend on two parameters (*λ′, μ′*), meaning only two out of the three birth–death parameters *λ, μ, ρ* can be inferred from the phylogenetic tree. Furthermore, the expressions for 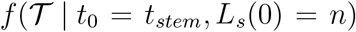 and 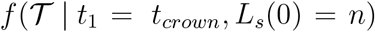 (i.e. the expressions where we condition on both the age of the process and the number of sampled tips) give the same result for *λ′, μ′* and for when the parameters are swapped to *μ′, λ′*. For complete sampling, [12] noticed this symmetry in 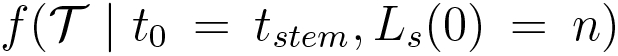 (This author mentioned that this special symmetry had also been independently observed by Monty Slatkin). Note that *μ*′ ≤ 0 is possible, whereas *λ*′ > 0, thus the switching is only well-defined if *μ*′ > 0.

## 5 Mapping from (*λ*′, *μ*′) to the birth–death model parameters (*λ, μ, ρ*) with consequences for maximum likelihood and Bayesian inference

When using the tree probability densities in a maximum likelihood inference framework, the expressions are maximized over the parameters for a given tree. Based on the five conditioned tree probability density equations, we should optimize over *λ*′ and *μ*′, with *λ*′ ∈ (0, ∞) and *μ*′ ∈ (−∞, ∞), instead of maximizing over the three parameters *λ, μ* and *ρ*, as the latter parameterization induces a ridge in the likelihood surface and thus optimization is problematic. This is equivalent to optimizing when assuming complete sampling (and allowing the ‘death rate’ *μ*′ to be negative) and, in a second step, assuming a sampling probability *ρ* and transforming from (*λ*′, *μ*′) to (*λ, μ*). We next investigate for which chosen values of *ρ* we can transform *λ*′, *μ*′ to *λ, μ*.

### Theorem 5.1.

*Let P denote the conditioned tree probability density for an arbitrary tree* 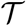 *given λ*′ ∈ (0, ∞) *and μ*′ ∈ (−∞, ∞). *The expression for P is given in the right column of Theorem 4.1. Each* (*λ*′, *μ*′) *has corresponding birth–death parameters* (*λ* ∈ (0, ∞), *μ* ∈ [0, 1)), *ρ* ∈ (0, 1]), *namely:*

- *Given μ*′ ≥ 0, *we obtain the same tree probability density P using the expression in the middle column of Theorem 4.1 with parameters* (*λ* = *λ*′/*ρ, μ* = *μ*′ – *λ*′ + *λ*′/*ρ*), *where ρ is any value in ρ* ∈ (0, 1].
- *Given μ*′ < 0, *we obtain the same tree probability density P using the expression in the middle column of Theorem 4.1 with parameters* (*λ* = *λ*′/*ρ*, *μ* = *μ*′ – *λ*′ + *λ*′/*ρ*), *where ρ is any value in* 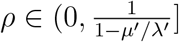.

Given the correlations among *λ, μ* and *ρ*, one may decide to perform a Bayesian Markov chain Monte Carlo analysis on *λ*′ ∈ (0, ∞), *μ*′ ∈ (–∞, ∞). Care has to be taken though regarding the priors, since these priors play out in non-straightforward ways. Assume, for example, that the analysis is performed by sampling *λ*′, *μ*′. For each sampled parameter pair, one might pick a *ρ* ∈ (0, 1] uniformly at random. Given that *μ*′ ≥ 0, this would yield a uniform distribution on the chosen *ρ*. However, given that some sampled parameter pairs reveal *μ*′ < 0, it follows that only a small *ρ*, namely 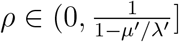 is possible, meaning that overall, the samples on *ρ* would be non-uniform, with a preference for small values of *ρ*. Thus, in the Bayesian setting, it is advantageous to estimate *λ, μ, ρ* in order to have control over their priors.

## 6 Mappings between birth–death model parameters (*λ, μ, ρ*) and 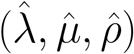

Next we characterize all birth–death parameters that are transformations of *λ, μ, ρ*.

### Theorem 6.1.

*Let* (*λ, μ, ρ*) *be birth–death parameters with the corresponding* (*λ*′, *μ*′). *There exists parameters* 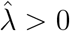, 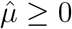, *and* 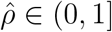 *with*

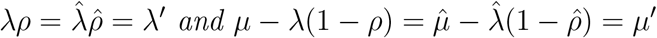

*if μ/λ* ≥ 1 *(for all* 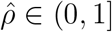) *and if μ/λ* < 1 *(for all* 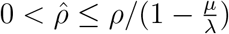).

Note that the parameters (*λ, μ, ρ*) and 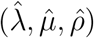 thus give thus rise to the same tree probability density.

### Corollary 6.2.

*With* 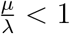 *(and thus* 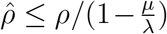*) a transformation always exists for* 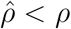. *However, a parameter transformation may not be possible for* 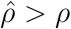 *(for example, if* 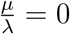, *we cannot transform to ρ*′ > *ρ)*.

Next we consider 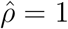 (i.e. the transformation to the case of complete sampling).

### Corollary 6.3.

*Let* (*λ, μ, ρ*) *be birth–death parameters with the corresponding* (*λ*′, *μ*′). *There exists a transformation to* 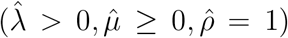 *if* 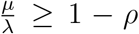. *If* 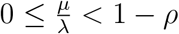, *no transformation exists*.

### 6.1 Consequences for the birth-death tree distribution

Sometimes, proofs of the properties of the conditioned tree distribution are carried out for complete sampling (i.e. for parameters 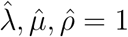). Such properties also hold for incomplete sampling if 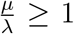 or if 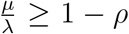. To include the parameter space, 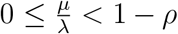 the proof needs to be done with explicitly acknowledging incomplete sampling. This was noticed already in [19].

### 6.2 Consequences for model choice regarding complete sampling

For a given phylogenetic tree, it is tempting to ask if a model with *ρ* = 1 or 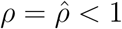 fits the data better. However, for every parameter combination (*λ, μ*, 1), we also find a parameter combination 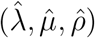 with both parameter triples having the same conditioned tree probability density. Moreover, there are parameter combinations 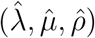 without a corresponding triplet where *ρ* =1 (see Corollary 6.2). Thus, the model with *ρ* < 1 always gets more support than the model with *ρ* = 1. In summary, such a test is meaningless because of the parameter correlations.

## 7 Discussion

Birth–death models have been studied for almost 100 years [22, 5]. However, surprising properties are still being uncovered. Here, we presented some unexpected symmetries in birth–death models, namely, fundamental birth–death probability distributions are invariant towards swapping the birth and the death rate. We explained this surprising observation in a special case, by using an argument that is both intuitive and precise, then derived more general symmetries algebraically. Second, we showed that a birth–death process with incomplete taxon sampling can be described phylogenetically through two parameters instead of three parameters due to parameter correlations, and that the two-parameter description again reveals symmetries.

Such correlations have important consequences for using birth–death models in phylogenetic and phylodynamic inference. In particular, the likelihood surface of the three birth–death parameters *λ, μ* and *ρ* for a given tree has a ridge casued by the correlations, and we can therefore only estimate two of the three parameters. Maximum likelihood estimation should thus be done over the two parameters. On the other hand, in Bayesian analysis, using the two-parameter description of the process would not allow us to use all prior information on the three original parameters and therefore using the original parameterization is advantageous.

Furthermore, we showed that for some of the parameter triplets (*λ, μ, ρ*), their two-parameter description is, in fact, equivalent to a birth–death process with complete sampling. However, in some cases, the resulting ‘death’ rate is negative, and thus the transformed parameters cannot always be considered as a birth–death process with complete-sampling. This means that we cannot simply prove properties of phylogenetic trees for complete sampling and then extrapolate to incomplete sampling, as we then miss some birth–death parameter combinations (namely the ones leading to a negative ‘death’ rate). Furthermore, testing whether the data are completely sampled (*ρ* = 1) or not (*ρ* < 1) is not informative, as the models with *ρ* < 1 always have more support: parameter triplets for incomplete sampling may only have corresponding complete sampling parameters with a negative ‘death’ rate, whereas birth and death rates under complete sampling have a corresponding triplet for all *ρ* ∈ (0, 1].

The birth–death model presented here is the simplest model for speciation and extinction, or for transmission and recovery. However, it has limitations for explaining the data, as it assumes exponential growth of the population, although populations cannot have unlimited growth, and it assumes that all individuals are dynamically equivalent. There has been considerable work on extending the birth-death model to address such limitations [8, 9, 16, 3, 17], but no symmetries and only very special parameter correlations have been observed [18]. It will be interesting to explore in the future whether the observed symmetries and correlations in our simple model are also present in these more complex models.

## 8 Acknowledgements

We wish to thank Joe Felsenstein and Nicolas Salamin for drawing our attention to the symmetry stated in Equation (1). We thank Bruce Rannala for pointing us to his work on tree symmetry in a special case [12]. TS is supported in part by the European Research Council under the Seventh Framework Programme of the European Commission (PhyPD: grant agreement number 335529).

## 9 Appendix: Proofs

### Proof of Theorem 3.1.

For *λ* > 0 and *ρ* > 0, the expressions are provided in [15], based on earlier work by [10, 21]. In fact, [15] requires *λ* > *μ*, but the proof is identical for *λ* < *μ*.

These expresions also hold for *λ* = 0 (and thus *μ* > 0). To see this, observe first that the probability *p*_1_(*t* | *λ* = 0, *μ, ρ*) is the product of the probability of no death *e*^−*μt*^ with the sampling probability *ρ*. Indeed this equation simplifies to:

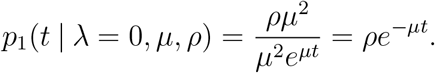

Second, notice that the probability *p_n_*(*t* | *λ* = 0, *μ, ρ*) for *n* > 1 is 0, as no birth event may occur. Indeed, by using the expressions above, we get 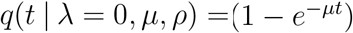 and thus *p_n_*(*t* | *λ* = 0, *μ, ρ*) = 0 for *n* > 1.

Finally, the probability *p*_0_(*t* | *λ* = 0, *μ, ρ*) is 1 − *p*_1_(*t* | *λ* = 0, *μ, ρ*) = 1 – *ρe*^−*μt*^. Again, the above equation simplifies:

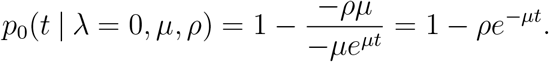

### Proof of Corollary 3.2.

Note that these equations can be derived from the supercritical case *λ* > *μ* by setting *λ* – *μ* = *ϵ* and using the property *e*^−*ϵ*^ ~ 1 – *ϵ* as *ϵ* → 0. In particular, for the expression in the denominator, we obtain:

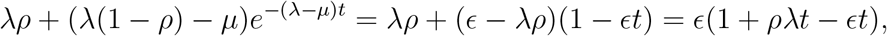

from which we directly get the expressions above.

Now we first prove Theorem 3.6 and then provide the proofs for the special case of complete sampling.

### Proof of Theorem 3.6.

The first three equations can be directly observed. For the last equation, observe that 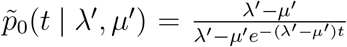. When swapping *λ*′ and *μ*′, we obtain:

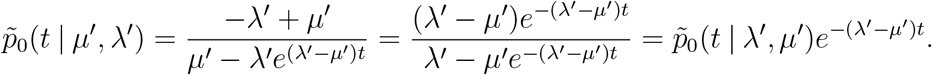

### Proof of Theorem 3.3.

For 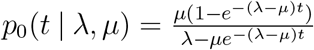, we obtain,

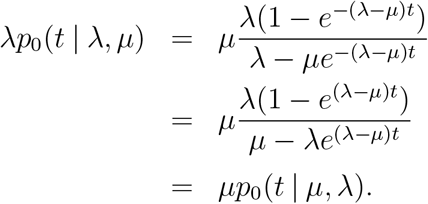

The remaining four equations are directly observed as special cases of Theorem 3.6. Alternatively, they can be established through simple algebraic rearrangements.

### Proof of Theorem 3.4.

First, we assume that both *λ* and *μ* are different from 0. For *m* = 0, we have *λ*^*n*^*p*_*n*,0_(*t* | *λ, μ*) = (*λp*_0_(*t* | *λ, μ*))^*n*^ = (*μp*_0_(*t* | *μ, λ*))^*n*^ = *μ^n^p*_*n*,0_(*t* | *μ, λ*) where the second equality follows from Thm. 3.3. For *m* > 0, we use a generating function argument. Let

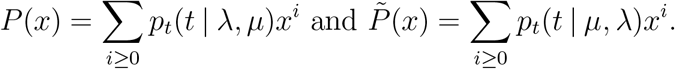

Theorem 3.3 gives:

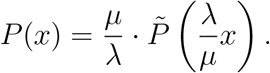

Now *p_n,m_*(*t* | *λ, μ*) is the coefficient of *x^m^* in *P*(*x*)^*n*^, which, by the previous equation, equals 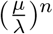 multiplied by the coefficient of *x^m^* in 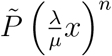. The latter coefficient is just *p_n,m_*(*t* | *λ, μ*) times 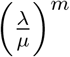. Thus 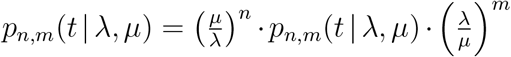, which leads to the claimed identities in the cases where *m* > 0 and *λ, μ* ≠ 0.

Finally, we prove the case where *λ*=0; the case where *μ* = 0 is then analogous. For the case *m* = *n, p_n,n_*(*t* | 0, *μ*) is the probability of no event happening within *t*, and thus *p_n,n_*(*t* | 0, *μ*) = *e*^−*μt*^ = *p_n,n_*(*t* | *μ*, 0). If *m* > *n*, *p_n,m_*(*t* | *λ, μ*) = 0 and thus the equation in the corollary is true. If *m* < *n*, then *p_n,m_*(*t* | *μ, λ*) = 0 and again the equation in the corollary is true.

### Proof of Theorem 5.1.

We can always transform *λ*′ ∈ (0, ∞) and *ρ* ∈ (0, 1] to *λ* ∈ (0, ∞) via *λ* = *λ*′/*ρ*. Second, since *μ*′ = *μ* – *λ*(1 – *ρ*) and *μ* ≥ 0, we have *μ* = *μ*′ – *λ*′ + *λ* = *μ*′ – *λ*′ + *λ*′/*ρ* ≥ 0, and thus *λ*′ – *μ*′ ≤ *λ*′/*ρ*. Thus, we can only transform *μ*′ to *μ* if this last inequality is fulfilled. This constrains our choices for *ρ* ∈ (0, 1]:

- For *λ*′ – *μ*′ > 0, the constraint is 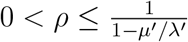.

– If *μ*′ ≥ 0, then 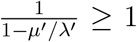, and thus there exists a transformation from *λ*′, *μ*′ to *λ, μ, ρ* for all *ρ* ∈ (0, 1].
– If *μ*′ < 0, we have 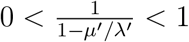 and thus we require 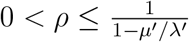 for a transformation to *λ, μ*.
- For *λ*′ – *μ*′ < 0 (implying *μ*′/*λ*′ > 1 and *μ*′ > 0), our constraint is, 1 ≥ *ρ* ≥ 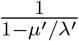. Since 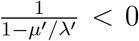, this means that a transformation for all *ρ* ∈ (0, 1] exists.
- For *λ*′ – *μ*′ = 0 (implying *μ*′ > 0), we require 0 ≤ *λ*′/*ρ* which is fulfilled for all *ρ* ∈ (0, 1].

### Proof of Theorem 6.1.

We can always transform *λ* ∈ (0, ∞) and *ρ* ∈ (0, 1] to 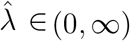 and 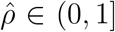 via 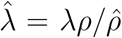. Second, since 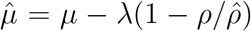 with 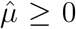, we need to determine for which 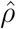 we have 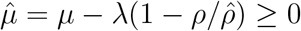.

- For *λ* = *μ*, we have 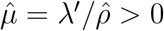 for all 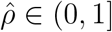.
- For *λ* ≠ *μ*, we obtain,

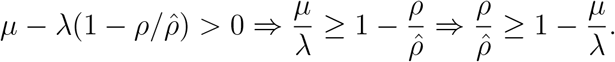

– For 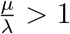, we have 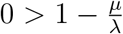 and thus we have 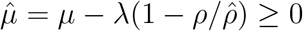 for all 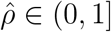.
– For 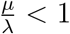, 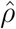 needs to fulfil 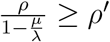 such that 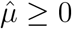.

## References

[1] David Aldous. Stochastic models and descriptive statistics for phylogenetic trees, from Yule to today. Stat. Sci., 16(1):23–34, 2001.

[2] David Aldous, Maxim Krikun, and Lea Popovic. Five statistical questions about the tree of life. Syst. Biol., 60(3):318–328, 2009.

[3] Rampal S Etienne, Bart Haegeman, Tanja Stadler, Tracy Aze, Paul N Pearson, Andy Purvis, and Albert B Phillimore. Diversity-dependence brings molecular phylogenies closer to agreement with the fossil record. Proc. Roy. Soc. Lond. B, 279(1732):1300–1309, 2012.

[4] J. Felsenstein. Inferring phylogenies. Sinauer Associates, Sunderland, Massachusetts, 8:8–5, 2004.

[5] David G. Kendall. On some modes of population growth leading to r. a. fisher’s logarithmic series distribution. Biometrika, 35(1/2):6–15, 1948.

[6] David G. Kendall. On the generalized “birth-and-death” process. Ann. Math. Statist., 19(1):1–15, 1948.

[7] A. Lambert and T. Stadler. Birth-death models and coalescent point processes: The shape and probability of reconstructed phylogenies. Theor. Pop. Biol., 90:113–128, 2013.

[8] W.P. Maddison. Estimating a binary character’s effect on speciation and extinction. Syst. Biol., 56(5):701–710, 2007.

[9] Hélène Morlon, Todd L Parsons, and Joshua B Plotkin. Reconciling molecular phylogenies with the fossil record. Proc. Natl. Acad. Sci. USA, 1081(39):6327–6332, 2011.

[10] S. C. Nee, R. M. May, and P.H. Harvey. The reconstructed evolutionary process. Phil. Trans. Roy. Soc. Ser B., 344:305–311, 1994.

[11] B. Rannala and Z. Yang. Probability distribution of molecular evolutionary trees: a new method of phylogenetic inference. J. Mol. Evol., 43:304–311, 1996.

[12] Bruce Rannala. Gene genealogy in a population of variable size. Heredity, 78(4):417, 1997.

[13] T Stadler. How can we improve accuracy of macroevolutionary rate estimates? Systematic biology, 62(2):321, 2013.

[14] Tanja Stadler. On incomplete sampling under birth-death models and connections to the sampling-based coalescent. J. Theor. Biol., 261(1):58–66, 2009.

[15] Tanja Stadler. Sampling-through-time in birth-death trees. J. Theor. Biol., 267(3):396–404, 2010.

[16] Tanja Stadler. Mammalian phylogeny reveals recent diversification rate shifts. Proc. Natl. Acad. Sci. USA, 108(15):6187–6192, 2011.

[17] Tanja Stadler and Sebastian Bonhoeffer. Uncovering epidemiological dynamics in heterogeneous host populations using phylogenetic methods. Phil. Trans. Roy. Soc. Ser. B, 368(1614):20120198, 2013.

[18] Tanja Stadler, Denise Kuhnert, Sebastian Bonhoeffer, and Alexei J Drummond. Birth-death skyline plot reveals temporal changes of epidemic spread in HIV and hepatitis C virus (HCV)). Proc. Natl. Acad. Sci. USA, 110(1):228–233, 2013.

[19] Tanja Stadler and Mike Steel. Distribution of branch lengths and phylogenetic diversity under homogeneous speciation models. J. Theor. Biol., 297:33–40, 2012.

[20] E. A. Thompson. Human evolutionary trees. Cambridge University Press, 1975.

[21] Z. Yang and B. Rannala. Bayesian phylogenetic inference using DNA sequences: A Markov chain Monte Carlo method. Mol. Biol. Evol., 17(7):717–724, 1997.

[22] G. U. Yule. A mathematical theory of evolution: based on the conclusions of Dr. J.C. Willis. Phil. Trans. Roy. Soc. Ser. B., 213:21–87, 1924.

